# Bacterial Zombies and Ghosts: Production of Inactivated Gram-Positive and Gram-Negative Species with Preserved Cellular Morphology and Cytoplasmic Content

**DOI:** 10.1101/458158

**Authors:** Rahwa Taddese, Clara Belzer, Steven Aalvink, Marien I. de Jonge, Iris D. Nagtegaal, Bas E. Dutilh, Annemarie Boleij

## Abstract

There are many approaches available to inactivate bacteria, each with a different efficacy, impact on cell integrity, and potential for application in high-throughput. The aim of this study was to compare these approaches and develop a standardized protocol for generation of intact Gram-positive as well as on Gram-negative “bacterial zombies”, i.e. cells that are metabolically dead with retained cellular integrity. Here, we introduce the term “bacterial zombies” in addition to “bacterial ghosts” to differentiate inactivated bacteria with preserved cellular integrity from those with perforated membranes, where DNA and cytoplasmic contents have been released. This differentiation of inactivated bacteria is important if the cell content is the subject of study, or if cell contents in the media may cause unwanted effects in downstream applications. We inactivated eight different bacterial species by treatment with beta-propiolactone, ethanol, formalin, sodium hydroxide, and pasteurization. Inactivation efficacy was determined by culturing, and cell wall integrity assessed by quantifying released DNA and visualization by scanning electron microscopy. Based on these results, we discuss the choice of bacterial inactivation methods, and conclude that beta-propiolactone and ethanol are the most promising approaches for standardized generation of bacterial zombies.

**IMPORTANCE:** For applications such as vaccination or analyses that are sensitive to bacterial growth, inactivated bacteria are preferred because they simplify the analyses and the interpretation of results. This study compared various bacterial inactivation treatments that maintain cell integrity and may be used in high-throughput. Our results demonstrated that beta-propiolactone and 70% ethanol were the best techniques to achieve these goals.

## INTRODUCTION

Bacterial inactivation refers to bactericidal methods that kill bacteria by damaging DNA or protein synthesis, resulting in termination of growth. While several techniques are available to achieve bacterial inactivation, most studies focus on single bacteria and there is no protocol for standardized and high-throughput inactivation of bacteria in general. Experiments that are sensitive to host-microbe interaction often involve inactivation of the bacteria before they are applied. Examples include immunization of humans and animals (vaccinations), analysis of cell response to bacterial outer-membrane structures, and the use of inactivated bacteria as carriers for drugs or antigens [1].

One challenge for a standardized protocol for bacterial inactivation lies in the diversity of the targeted bacteria. For example, the structurally different walls of Gram-positive and Gram-negative bacteria may inhibit certain inactivation treatments. One example is the production of bacterial ghosts by using a plasmid with an E gene insert derived from bacteriophage ФX174, that lyses Gram-negative bacteria by forming a lysis tunnel across the double membrane, where the DNA and cytoplasmic contents escape [1]. Additionally, DNA is then degraded by beta-propiolactone (BPL) and/or staphylococcal nuclease A. Finally, inactivated bacteria are lyophilized to ensure inactivation. This procedure inactivates a range of Gram-negative bacteria, but it is not compatible with Gram-positive bacteria. A double membrane such as what Gram-negative bacteria have is necessary for the formation of a lysis tunnel [1; 2].

A second challenge lies in finding methods for application in high-throughput. Gram-positive bacterial ghosts were created for *Listeria monocytogenes* (referred to as *L. monocytogenes* ghosts or LMGs) using a combination of chemicals in a series of steps [3]. With these chemicals, the minimum inhibitory concentrations had to be determined beforehand to produce inactivated bacteria with preserved structure. The minimum inhibitory concentrations may differ for individual bacteria, making certain inactivation treatments unsuitable for high-throughput protocols.

A third challenge when standardizing the inactivation protocol lies in the methods to quantify bacterial cells. Several protocols require that comparable amounts of bacteria are available to ensure efficient inactivation. There are numerous methods to quantify bacteria with different accuracy and different suitability for application in high-throughput, including plating and counting colony forming units (CFU), 4,6-diamidino-2-phenylindole (DAPI)-staining, fluorescent-activated cell sorting (FACS), and measuring optical density (OD) [4; 5]. DAPI and FACS require further processing of bacteria which end up being time-consuming. Measuring OD may be the quicker and easier option, albeit less accurate than plating and counting CFUs. Therefore, next to the plating, OD may be measured to proceed with experiments immediately while incubating bacteria on the plates.

“Bacterial ghosts” are inactivated bacteria whose cytoplasmic content including DNA have been lost via perforated membranes [1; 3]. Here, we introduce the term bacterial zombies to indicate intact inactivated bacteria whose DNA and cytoplasmic content remains within the cell. We compared five protocols for their ability to effectively inactivate bacteria while preserving the bacterial membrane structure, applicability to different bacterial strains, and the possibility for standardization, including four chemical treatments beta-propiolactone (BPL), ethanol, formaldehyde (further referred to as formalin), sodium hydroxide (NaOH), and one physical treatment, i.e. pasteurization.

BPL is commonly used for the inactivation of viruses for vaccinations. It acts mainly by damaging the DNA [6]. Previous research has found that BPL chemically modifies membrane fusion proteins and antigen proteins on the surface of Influenza viruses whereas another study showed that protein structures and foldings are not greatly affected in rabies viruses [7; 8; 9]. BPL has been previously applied on bacterial cells, but it is not known if BPL affects the bacterial surface structure.

Ethanol is generally used for cleaning surfaces and sterilization due to its bactericidal effects. It inactivates bacteria by disrupting the cell membrane, dehydrating the bacterial cells and denaturing proteins. Interestingly, spores are not inactivated by ethanol [10; 11].

Formalin is applied as fixating agents for histological tissue samples. It also acts on bacteria by interacting with nucleic acids and proteins and by dehydrating the cells while keeping the structure intact [11]. However, cross-links with proteins could change epitopes for antibodies on the cell surface [12].

NaOH has been previously used to create bacterial ghosts. The minimum inhibitory concentration of NaOH was determined to create bacterial ghosts from *Staphylococcus aureus* [13]. Using scanning electron microscopy (SEM), they demonstrated that NaOH perforates the bacterial membrane where the bacterial DNA escapes and is degraded. Except for the pores, the membrane and cell wall of the bacteria were shown to remain intact.

Pasteurization is a commonly used method in the food industry to extend shelf-life of products without destroying essential nutrients. Common bacterial pasteurization works by raising the temperature up to 70°C for a maximum of 30 minutes, denaturing proteins and cell membranes of bacteria [14].

The techniques mentioned above demonstrate effective bacterial inactivation, but it is mostly unknown whether the surface structures are preserved for both Gram-positive and Gram-negative bacteria. The aim of this study is to evaluate the protocols for the efficacy of bacterial inactivation while preserving the surface structure and for their potential to use them in high-throughput inactivation.

## MATERIALS AND METHODS

### Bacterial strains and growth conditions

Eight strains of bacteria from five phyla were selected as representatives for the structural differences in bacterial cell walls of Gram-positive and Gram-negative strains to evaluate the inactivation efficacy of each method (Table 1). The anaerobic bacteria *Bacteroides fragilis* 9343 NTBF, *Parabacteroides distasonis* 3999B T(B)4, *Fusobacterium nucleatum* patient isolate NTB17 (from the Radboudumc strain collection), *Akkermansia muciniphila* ATCC BAA-835 were cultured in an anaerobic jar using sachets (Thermo Fisher Scientific, USA) in Brain-Heart-Infusion (BHI) broth (Sigma-Aldrich, USA) supplemented with L-cysteine (Sigma-Aldrich, USA), yeast extract (BD, USA), hemin (Sigma-Aldrich, USA), vitamin K1 (Sigma-Aldrich, USA) for 48 hours at 37°C. Facultative aerobic strains such as *Salmonella enteric* serovar Typhimurium NTB6 [15] (further designated as *S. typhimurium*), *Streptococcus gallolyticus* subsp. *gallolyticus* UCN34 (further designated as *S. gallolyticus*), *Escherichia coli* NC101 delta pks and *Lactococcus lactis* IL1403 were cultured aerobically in BHI broth overnight at 37°C with 5% CO_2_.

**Table 1:** Strains, their phyla and properties.

| Bacteria | Phyla | Oxygen requirements | Gram stain |
| --- | --- | --- | --- |
| *Akkermansia muciniphila* | Verrucomicrobia | Anaerobe | Negative |
| *Bacteroides fragilis* | Bacteroidetes | Anaerobe | Negative |
| *Fusobacterium nucleatum* | Fusobacteria | Anaerobe | Negative |
| *Parabacteroides distasonis* | Bacteroidetes | Anaerobe | Negative |
| *Escherichia coli* | Proteobacteria | Facultative aerobe | Negative |
| *Lactococcus lactis* | Firmicutes | Facultative aerobe | Positive |
| *Streptococcus gallolyticus* | Firmicutes | Facultative aerobe | Positive |
| *Salmonella typhimurium* | Proteobacteria | Facultative aerobe | Negative |

Following incubation, optical density at 620 nm was measured in the microplate reader Infinite F50 (Tecan, Switzerland) and samples were centrifuged at maximum speed (16,100 rcf) for 10 minutes. Supernatants were discarded and OD_620_ was adjusted to 1.0 in 0.9% sodium chloride buffer (B Braun Melsungen AG, Germany) for treatments with BPL (Acros Organics, Thermo Fisher Scientific, USA) and pasteurization. For NaOH, ethanol (all from Merck, Germany) and formalin (formaldehyde solution about 37%, Merck, Germany) treatments, pellets were resuspended at OD_620_ of 1.0 in NaOH, ethanol and formalin, respectively.

### Inactivation treatments

***BPL treatment*** Bacteria were dissolved in 0.9% sodium chloride buffer. BPL was added to the buffer at concentration of 1:2,000 (v/v) and samples were incubated rotating at 150 rpm at 4°C overnight (modified protocol from *Gonçalves* et al. [16]). After incubation, samples were placed at 37°C for 2 hours to inactivate BPL. No rinsing was required. ***NaOH, ethanol and formalin treatments*** Bacterial pellets (Table 1) were resuspended in 6 mg/mL NaOH solution, 70% ethanol, and formalin, and incubated at room temperature for 5 minutes. ***Pasteurization*** Bacteria were heat-treated in a dry block heater (Grant, UK) at 70°C for 30 minutes. Hereafter, samples were immediately placed on ice.

Immediately after incubation, 15 μL of each treatment was transferred into the first serial dilutions to obtain colony forming units. Bacteria were then centrifuged (16,100 rcf, 10 minutes) where the top part of the supernatants were transferred into a new eppendorf tube to measure the DNA concentration. Using 0.9% sodium chloride buffer, bacteria were resuspended for rinsing. This was repeated by centrifuging and resuspending bacteria in the same buffer again.

### Colony forming unit (CFU) determination

Following inactivation treatments, 10-fold serial dilutions were plated with treated and untreated bacteria to determine the CFU and compare the inactivation efficacy. Three drops (15 μL each) from each dilution were placed on BHI agar plates and incubated aerobically (37°C, 5% CO_2_, overnight) and anaerobically (37°C, 48 hours) for facultative aerobes and anaerobes, respectively. After incubation, colonies were counted and CFU/mL were calculated for each inactivation method. The CFU/mL also gives information on the amount of bacteria prior to treatment at OD_620_ of 1.

### Measurement of the concentration of extracellular DNA

After treatments, bacteria were centrifuged at 16,100 rcf for 10 minutes and DNA concentration was measured from the top fraction of the supernatants at 260 nm using NanoDrop ND-1000 (Isolagen Technologies, USA).

### Scanning electron microscopy (SEM)

***Fixation*** Following DNA measurements, remaining supernatants were completely removed and bacterial pellets were fixed in 2% glutaraldehyde diluted in 0.1 M cacodylate buffer overnight at 4°C. Samples were centrifuged, washed with 0.1 M cacodylate buffer and stored at 4°C in same buffer until use. When ready for SEM visualization, post-fixation was performed. For this, buffer was removed by centrifugation and bacteria were incubated in 1% osmium tetroxide for 1 hour at room temperature. Samples were centrifuged and washed with deionised water. ***Dehydration*** Bacteria were placed on filter papers with pore size of 5 to 13μm (Thermo Fisher Scientific, USA) that were cut to approximately 12 mm in diameter and hydrated with few drops of water. Vacuum-suction was applied for bacteria to stick on the filters. Dehydration was performed in ethanol series of 50%, 70%, 80%, 96% and 100%. Filters must be kept wet at all times, so some ethanol was always left at the bottom every time the solution was exchanged. ***Drying*** Hexamethyldisilazane (HMDS, Sigma-Aldrich, USA) was used for drying bacterial samples. For drying the filters, three HMDS dilutions (2:1, 1:1 and 1:2) were prepared using 100% ethanol. Also here, filters must be kept wet by leaving some solution during exchange. ***Coating*** Filters were attached to aluminum Zeiss pin stubs with 12.7 mm diameter (MicrotoNano, The Netherlands) and coated with gold in HHV Scancoat Six bench-top sputter coater (HHV Ltd., UK) 3 times for 30 seconds each. ***SEM visualization*** Preservation and damage of bacterial surface structures of the bacteria after inactivation treatments were compared to that of untreated bacteria at comparable magnifications and settings using Zeiss Sigma 300 (Carl Zeiss, Germany).

### Quantification of dents and ECS in SEM pictures

For quantification, pictures with a maximum of 50 bacteria were selected. The number of dents and ECS on each cell were counted and the percentages of total dents or ECS were calculated.

### Fluorescein isothiocyanate (FITC) and 4′,6-diamidino-2-phenylindole (DAPI) labeling

***FITC and DAPI labeling of live bacteria*** Bacteria were grown overnight and OD_620_ was adjusted to 1.0. Bacteria were washed once with phosphate buffered saline (PBS) and centrifuged at 16,100 rcf for 3 minutes. During centrifugation, FITC (Sigma-Aldrich, USA) was dissolved in dimethyl sulfoxide (DMSO, PanReac AppliChem) at 5 mg/mL and diluted in PBS at 0.5 mg/mL (FITC/PBS mixture). For each bacterium negative controls were live bacteria, stained with FITC but not DAPI (Prolong^Tm^ Gold Antifade Mountant with DAPI, Thermo Fisher Scientific, USA) (see Table 2). Positive controls are stained with both, DAPI and FITC. Following centrifugation, bacterial pellets and positive controls were resuspended in 1 mL FITC/PBS mixture. Bacteria were incubated rotating in the dark for 30 minutes for labeling. Afterwards, they were washed with PBS 3 times to remove non-bound FITC (centrifugations at 16,100 rcf for 3 minutes). ***Inactivation treatments*** Except for positive and negative controls, bacteria were inactivated with various treatments. ***Fixation*** Inactivated cells were centrifuged at 16,100 rcf for 3 minutes and pellets were resuspended in formalin for 5 minutes. Formalin was washed out with PBS 3 times (centrifugations at 16,100 rcf for 3 minutes). ***Staining with DAPI and preparation for microscopy*** After washing out formalin, bacteria were dissolved in PBS. 5 μL of bacteria were placed on slides at a dilution of 10^1^ or 10^2^. Bacteria were air dried on the slides and then covered with 1-2 drops of Prolong^Tm^ Gold Antifade Mountant with DAPI and cover slips. For negative controls, Quick-D mounting medium (Klinipath, The Netherlands) was used. Finally, bacteria were visualized under the fluorescence microscope at 100x magnification (with oil). Slides were stored in the fridge at 4°C in the dark until imaging for a maximum of 3 months.

**Table 2:** Overview of steps in FITC experiment.

|  | Controls |  | Each Treatment<br>(Ethanol, formalin,<br>pasteurization, NaOH) |
| --- | --- | --- | --- |
|  | Positive control | Negative control | Fixed |
| 1. FITC staining | √ | √ | √ |
| 2. Fixation | √ | √ | √ |
| 3. DAPI staining | √ | X | √ |
| 4. Visualization | 10<sup>1</sup> and 10<sup>2</sup> | 10<sup>1</sup> and 10<sup>2</sup> | 10<sup>1</sup> and 10<sup>2</sup> |

## RESULTS

The efficacy of the inactivation treatments; BPL, ethanol, formalin, NaOH, and pasteurization was examined, aiming to obtain intact bacterial zombies. For this, eight anaerobic and facultative aerobic bacterial strains from five phyla were selected based on their differences in cell wall structures, including two Gram-positive and six Gram-negative strains (see Table 1).

### Efficacy of inactivation

To assess the efficacy of inactivation treatments, CFU/mL prior (at OD_620_ of 1.0) and post treatments were calculated by counting colonies from agar plates (see Supplementary Table S1). The inactivation efficacy of BPL, ethanol, formalin, and NaOH was 100%, with no bacterial colonies observed on the plates. While 30 minutes of pasteurization inactivated >99.65% bacteria for all strains, Figure 1 shows that some cells of *E. coli*, *S. typhimurium*, *P. distasonis*, *L. lactis* and *A. muciniphila* survived the pasteurization process, rendering this inactivation approach the least efficacious of all those tested.

**Figure 1:**
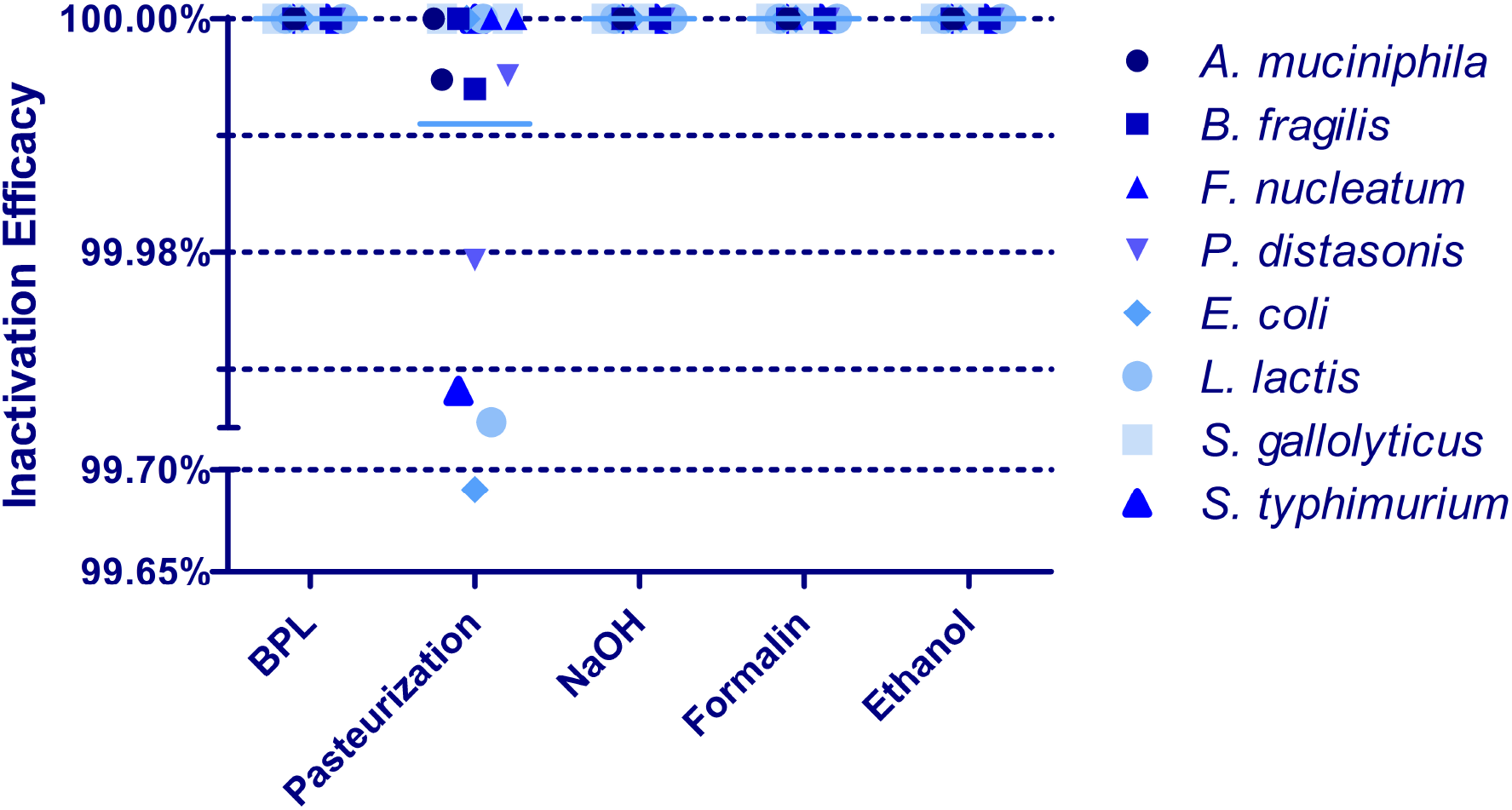
Efficacy of inactivation treatments for individual bacteria. Complete inactivation of bacteria, i.e. 0 CFU/mL, was taken as 100%. Mean values calculated across all bacteria (displayed by horizontal blue line): Pasteurization 99.97±0.07%, and 100% for untreated bacteria, BPL, NaOH, Formalin and Ethanol. Pasteurization displayed colonies for *E. coli*, *S. typhimurium*, *P. distasonis*, *L. lactis* and *A. muciniphila* compared to untreated control.

### Maintenance of cellular integrity: Quantification of released extracellular DNA in supernatants

As defined above, bacterial zombies differ from bacterial ghosts in that they have retained their cellular integrity. We applied several complementary methods to evaluate the effects of the various inactivation treatments on cellular integrity of inactivated bacteria. We measured the release of extracellular DNA (exDNA) in the supernatants, imaged the bacteria with SEM, labeled surface proteins with FITC, and labeled intracellular DNA with DAPI.

Figure 2 reveals significant differences in exDNA of individual bacteria between the different inactivation methods. Supernatants of untreated bacteria (served as negative control) generally contained low amounts of exDNA (mean ± standard deviation, 30.67 ± 17.45 ng/μL). Striking was the high exDNA (116.4 ± 33.33 ng/μL) released in all NaOH-treated samples when compared to the negative control. ExDNA in BPL-treated bacteria was consistently low. Here, the mean value (29.74 ± 11.34 ng/μL) was similar to the negative control (30.67 ± 17.45 ng/μL). When comparing pasteurization, ethanol and formalin with each other, the standard deviations of pasteurization (60.96 ± 18.85 ng/μL) and ethanol (42.50 ± 18.14 ng/μL) were smaller than that of formalin (57.88 ± 56.94 ng/μL). From all the treatments, the mean released exDNA concentrations of ethanol (42.50 ± 18.14 ng/μL) and BPL (29.74 ± 11.34 ng/μL) were closer to that of the negative control (30.67 ± 17.45 ng/μL), and all their mean concentrations were under 50 ng/μL. The results of quantified exDNA in supernatants indicate that BPL and ethanol inactivate bacteria while keeping the cells intact, creating bacterial zombies. Pasteurization, NaOH and formalin break down cell walls releasing the DNA and generating bacterial ghosts.

**Figure 2:**
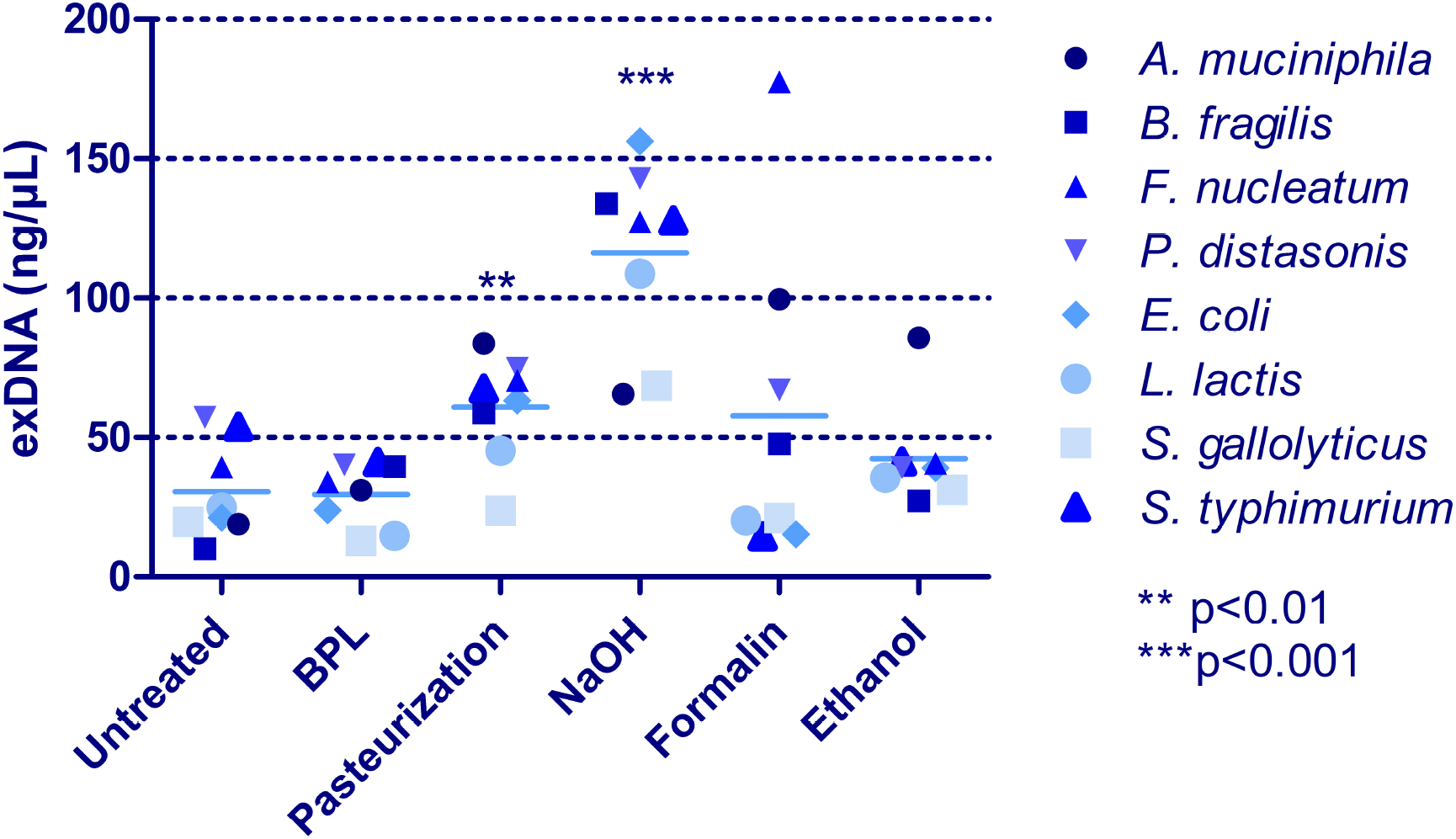
Extracellular DNA (exDNA) of supernatants from inactivated and untreated individual bacteria. Values were measured by Nanodrop, of which the mean was calculated across all bacteria (displayed by horizontal blue line): Untreated bacteria 30.67±17.45 ng/μL, BPL 29.74±11.34 ng/μL, Pasteurization 60.96±18.85 ng/μL, NaOH 116.4±33.33 ng/μL, Formalin 57.88±56.94 ng/μL, Ethanol 42.50±18.14 ng/μL. NaOH and pasteurization displayed significantly higher exDNA release than untreated control.

Interesting was also that the exDNA values of Gram-positive bacteria was below the average with all bacteria that were treated and untreated. This indicates that the cell walls of Gram-positive bacteria may be more stable and not easily disrupted using the various inactivation treatment procedures.

### Maintenance of cellular integrity: Imaging of inactivated bacteria by scanning electron microscopy

To visualize whether surface structures of inactivated bacteria were intact or damaged, SEM imaging was carried out.

We investigated the BPL and ethanol treatments using SEM imaging because they had 100% inactivation efficacy and little leakage of DNA. Since NaOH had a relatively high exDNA in most bacterial supernatants (Figure 2), we included this treatment as a positive control of a disrupted cell wall structure. The Gram-positive facultative aerobe *S. gallolyticus* and the Gram-negative anaerobe *A. muciniphila* were selected for visualization with SEM because their exDNA release was consistently below 100 ng/μL with all treatments. The effects of inactivation treatments on the surface structures and shapes of treated and untreated cells were compared (Figure 3).

**Figure 3:**
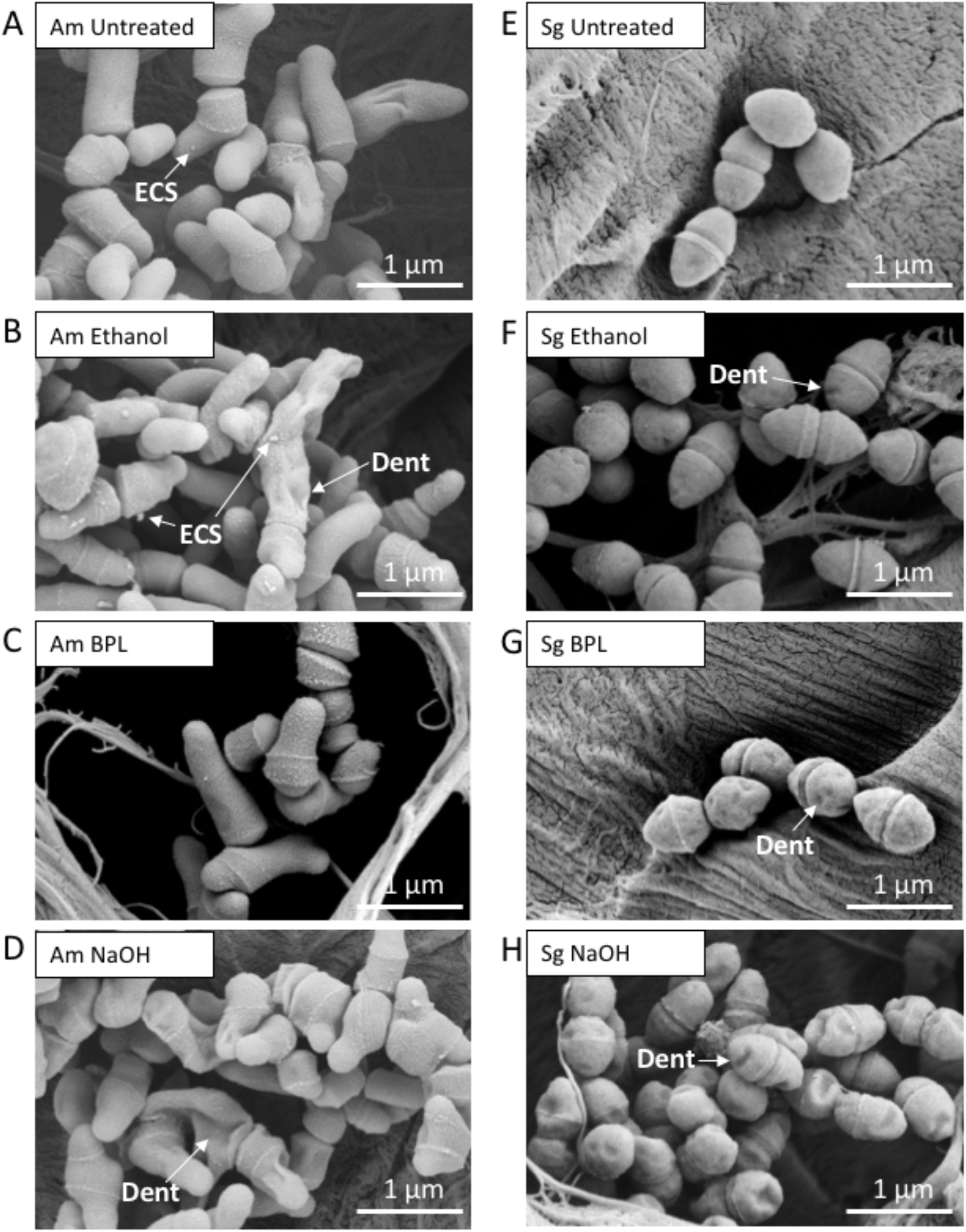
SEM images of untreated and inactivated *A. muciniphila* (Am) and *S. gallolyticus* (Sg). Shown are untreated Am (A) and Sg (E), ethanol-treated Am (B) and Sg (F), BPL-treated Am (C) and Sg (G), and NaOH-treated Am (D) and Sg (H). Dent-arrows point at damages on treated bacteria. Extracellular structures (ECS)-arrows points at extracellular structures on Am.

As shown in Figures 3A and 3E, *A. muciniphila* grows in long rods whereas *S. gallolyticus* forms diplococci. Normal untreated as well as treated bacteria clearly show the division ring. Ethanol-treated *A. muciniphila* exhibit increased dents and extracellular surface structures (ECS, arrows in Figure 3B) but most bacteria seem similar to untreated cells (Figure 3A). *S. gallolyticus* also shows dents in the outer structure but the shape of bacterial cells is preserved (Figure 3F). Figures 3C and 3G reveal that the structures of BPL-treated *A. muciniphila* and *S. gallolyticus* are preserved. With *A. muciniphila* there is minimal structural damage observed. *S. gallolyticus* appears to have more dents than ethanol-treated *S. gallolyticus* (Figure 3F). Hence, BPL and 70% ethanol are demonstrated to generate inactivated intact bacteria (Figures 3B-C, 3F-G). As expected, the NaOH-treated *A. muciniphila* and *S. gallolyticus* bacteria displayed the most severe structural damage, even though they had the lowest exDNA release with NaOH as measured using Nanodrop (Figure 2). Studies have shown that using NaOH perforates the bacterial cell wall, allowing the DNA and the cytoplasmic contents to spill out of the cell [3; 13]. Our SEM pictures (Figures 3D and 3H) indicate the creation of structurally preserved inactivated *A. muciniphila* and *S. gallolyticus* bacteria when using a very low NaOH concentration of 6 mg/mL.

Damage in SEM images was quantified (Tables 3 and 4) by assessing the percentages of cells with dents and ECS. The controls exhibit some dents which might be due to preparations for SEM imaging. Interestingly, the treatment NaOH seems to have caused most damage in both bacteria (27.6% for Am, 64.9% for Sg) compared to the controls (9.1% for Am, 27.6 for Sg). There are dents in all images except for BPL-treated *A. muciniphila* and untreated *S. gallolyticus*, where NaOH presented most dents in both bacterial species. As for the ECS, it looks to be specific for *A. muciniphila* since there was only 1 out of 81 *Streptococci* observed with ECS. *Akkermansia* showed most ECS with the ethanol treatment. Further studies are required to examine the ECS in *A. muciniphila*.

**Table 3:** Quantification of dents in SEM pictures.

|  | Number of cells with # dents |  |  |  |  |  | Total cells | % of cells with dents |
| --- | --- | --- | --- | --- | --- | --- | --- | --- |
|  | 0 dents | 1 dents | 2 dents | 3 dents | 4 dents | ≥ 5 dents |  |  |
| **Am Untreated** | 20 | 2 | - | - | - | - | 22 | 9.1 |
| **Am Ethanol** | 20 | 2 | - | - | - | 1 | 23 | 13.0 |
| **Am BPL** | 12 | - | - | - | - | - | 12 | 0 |
| **Am NaOH** | 21 | 4 | 1 | 1 | 2 | - | 29 | 27.6 |
| **Sg Untreated** | 6 | - | - | - | - | - | 6 | 0 |
| **Sg Ethanol** | 21 | 4 | 4 | 1 | - | - | 30 | 30 |
| **Sg BPL** | 4 | 1 | - | 2 | - | 1 | 8 | 50 |
| **Sg NaOH** | 14 | 14 | 9 | 1 | - | - | 37 | 64.9 |

**Table 4:** Quantification of ECS in SEM pictures.

|  | Number of cells with # ECS |  |  |  |  | Total cells | % of cells with ECS |
| --- | --- | --- | --- | --- | --- | --- | --- |
|  | 0 ECS | 1 ECS | 2 ECS | 3 ECS | ≥ 4 ECS |  |  |
| **Am Untreated** | 21 | 1 | - | - | - | 22 | 4.5 |
| **Am Ethanol** | 16 | 5 | 1 | 1 | - | 23 | 30.4 |
| **Am BPL** | 11 | 1 | - | - | - | 12 | 8.3 |
| **Am NaOH** | 28 | 1 | - | - | - | 29 | 3.4 |
| **Sg Untreated** | 6 | - | - | - | - | 6 | 0 |
| **Sg Ethanol** | 30 | - | - | - | - | 30 | 0 |
| **Sg BPL** | 8 | - | - | - | - | 8 | 0 |
| **Sg NaOH** | 36 | 1 | - | - | - | 37 | 2.7 |

### FITC and DAPI labeling

FITC and DAPI labeling was performed to examine the membrane integrity of *S. gallolyticus* (Figure 4). FITC binds to outer surface proteins while DAPI binds to DNA. Positive controls were labeled with both, FITC and DAPI to demonstrate functioning of both stains. Negative controls were only stained with FITC but not DAPI to ensure that FITC stain does not appears on DAPI channel.

**Figure 4:**
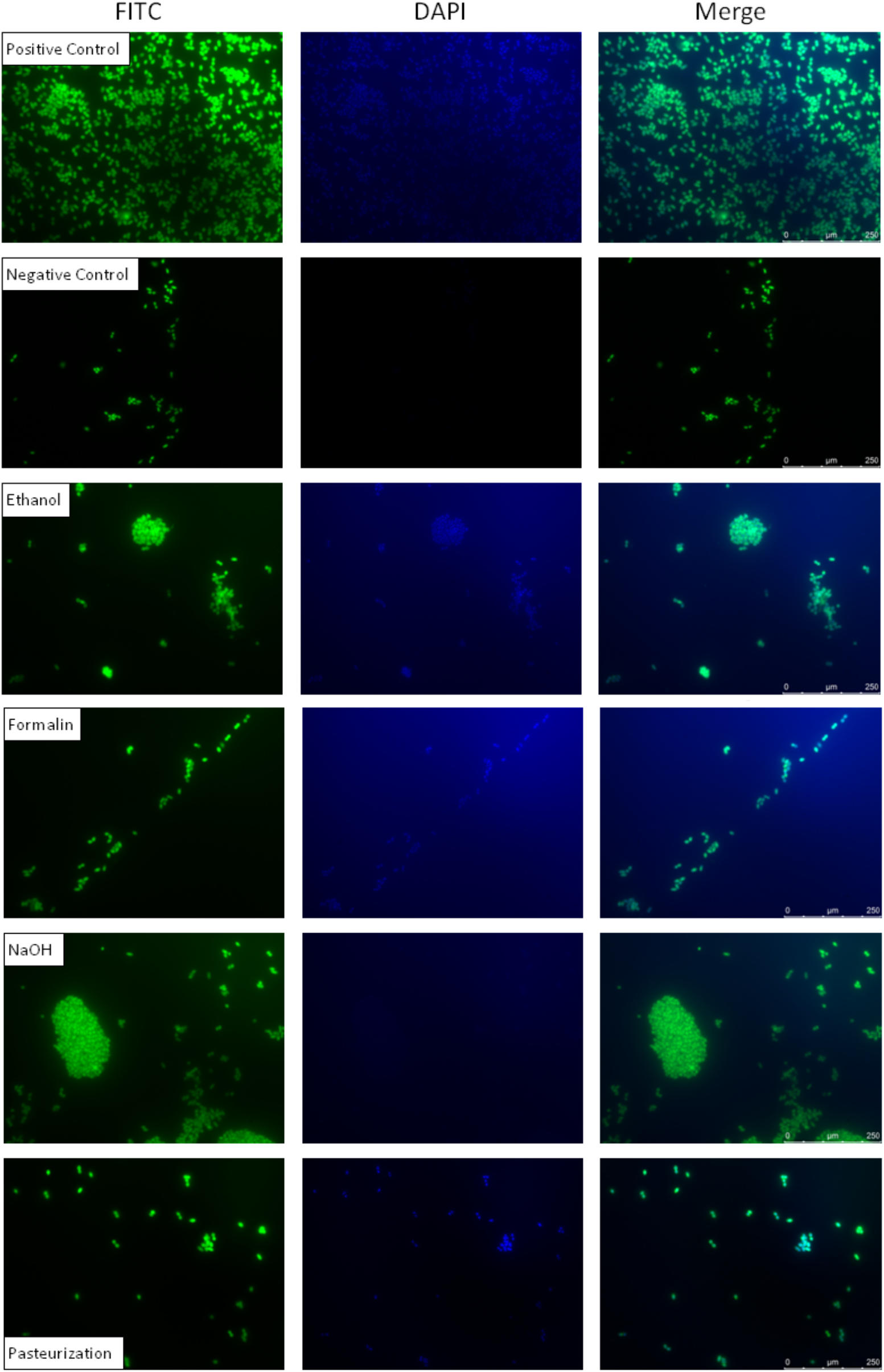
FITC and DAPI labeling of untreated and inactivated S. gallolyticus. The first and second rows are positive and negative control (only FITC labeled), respectively. The next rows show S. gallolyticus treated with ethanol, formalin, NaOH and pasteurization, and labeled with FITC and DAPI. Note that no DAPI stain was used in negative control.

In Figure 4, both controls with FITC showed intensely colored, intact cells. Treated bacteria also seem to be intact since the green color is intense and comparable to both controls. DAPI specifically stained the DNA in positive control, verifying the presence of DNA within intact cells. As expected, bacteria in negative control were not DAPI stained so no staining is visible. DAPI is intensely labeled in bacteria treated with ethanol, formalin, and pasteurization which confirms that the DNA was retained in the nucleus. In contrast, NaOH did not exhibit any staining, indicating that DNA escaped the bacterial cells. To conclude, while all inactivation treatments preserved the cell structure (FITC staining), NaOH inflicted considerable damage to the membrane integrity, releasing the DNA (DAPI staining).

## DISCUSSION

We aimed to examine different methods for inactivation of bacteria while preserving the membrane integrity, i.e. creating “bacterial zombies”. We chose eight bacteria from five phyla with different Gram stains and oxygen requirements to inactivate with BPL, pasteurization, ethanol, formalin, and NaOH. Our results demonstrate that BPL and ethanol showed 100% inactivation efficacy and minimal leakage of DNA. These methods may be applied in a standardized high-throughput inactivation protocol.

The techniques formalin and NaOH were not suitable for a protocol where preserved structure is required because they did not fulfil the aforementioned criteria of minimal DNA leakage. While pasteurization can be easily performed, not all bacteria were inactivated in our experiments in contrast to previous reports [14]. This difference might be the result of a different experimental setup, e.g. we used a heat block instead of a water bath and a larger volume for pasteurization, possibly interfering with the conductivity. Perhaps the inactivation efficacy might be improved by increasing the temperature and/or the incubation time, but this was not further tested.

Furthermore, NaOH treatment inactivated all bacteria and the protocol was simple to carry out, but it had highest DNA release and surface structures exhibited many dents. Therefore, we suggest that NaOH is a good technique for generating “bacterial ghosts” whose cellular content is released [1; 3].

With BPL, all tested bacteria were inactivated. A major advantage of the BPL treatment is that the protocol does not include a washing step after inactivation, so there is minimal loss of bacterial material. Thus, BPL inactivation can be used when dealing with low-input samples. The disadvantage is that BPL must be handled very carefully when conducting experiments. Mice experiments showed that BPL lead to formation of carcinoma upon skin exposure and nasal cancer upon inhalation [17; 18; 19]. Thus, BPL should be used only under an air replacement hood. After bacterial inactivation, BPL-treated samples are placed at 37°C for 2 hours to inactivate BPL. Then the samples are safe to use for further applications.

Ethanol and formalin completely inactivated all tested microbes within 5 minutes. Both these protocols were readily executed and may be performed in high-throughput. The ethanol treatment did not show exDNA concentrations higher than untreated samples. Note that here; the absence of exDNA is not the result of DNA precipitation by ethanol, since no resuspended DNA could be detected, even after centrifugation of any potential precipitate (Supplementary Figure S1). Moreover, DAPI staining of ethanol treated cells showed that the DNA remained concentrated inside the cells (Figure 4). A disadvantage of ethanol may be the washing out of lipoproteins from the cell-surface, so this approach might be less suitable for antibody production or immunization [20; 21].

ExDNA release of formalin was widely scattered (Figure 2). This may indicate that formalin can disrupt the structures of certain bacteria more than others, potentially making this treatment unreliable for generalized application.

## Conclusion

This study for the first time compares different methods for bacterial inactivation, on a range of microbiota members, with an outlook to their potential use in high-throughput. Our results suggest that inactivation treatments with BPL or ethanol are best for standardized high-throughput inactivation and creation of bacterial zombies with intact surface structures.

## LIST OF NON-STANDARD ABBREVIATIONS

BHI: brain-heart-infusion
BPL: beta-propiolactone
CFU: colony forming units
DAPI: 4′,6-diamidino-2-phenylindole
ECS: extracellular structures
FACS: fluorescent-activated cell sorting
FITC: fluorescein isothiocyanate
HMDS: hexamethyldisilazane
NaOH: sodium hydroxide
OD: optical density
PBS: phosphate buffered saline
SEM: scanning electron microscopy

## ACKNOWLEDGEMENTS

RT was supported by Radboud Institute for Molecular Life Sciences (RIMLS 014-058). BED was supported by the Netherlands Organization for Scientific Research (NWO) Vidi grant 864.14.004. AB was supported by the Netherlands Organization for Scientific Research (NWO) Veni grant 016.166.089.

## AUTHOR CONTRIBUTIONS

RT, BED and AB contributed conception and design of the study. RT performed experiments and wrote the first draft of the manuscript. All authors supervised and discussed experiments, contributed to manuscript revision, read and approved the submitted version.

